# The case for species selection

**DOI:** 10.1101/084046

**Authors:** Carl Simpson

## Abstract

The mere existence of speciation and extinction make macro-evolutionary processes possible. Speciation and extinction introduce discontinuities in the microevolutionary change within lineages by initiating, disrupting, and terminating the continuity of species lineages. Within a clade, speciation and extinction become potent means of macroevolution in and of themselves. This process, termed species selection, is a macroevolutionary analogue of natural selection, with species playing an analogous part akin to that played by organisms in microevolution. That said, it has proven difficult to think about levels of selection. The concept of species sorting was introduced to help our thinking on this issue by identifying two aspects inherent in hierarchical systems can confuse our attempts to understand them: uncertainty in the level that selection acts and uncertainty about if the pattern of selection is in fact caused at all. Thanks to insights from evolutionary transitions in individuality, we now know more about how to identify the level of selection and how to parse the causal structure in hierarchical evolutionary circumstances. We know that if the fitness of organisms causes the fitness of more inclusive species then they must covary. However, there is no evidence of such a covariance between fitnesses at these two levels. This covariance is just not observed; neither between cells and organisms nor between organisms and species. Rather, speciation and extinction rates appear to be completely divorced from organismal fitness. With this insight, the concept of species sorting shrinks so that it only covers the two processes of species selection and drift. I argue that we are better off focusing on understanding the processes of species selection and drift and that there is therefore no further need for the concept of species sorting.

## How are trends produced?

Macroevolution as a field of study, took shape in the 1970s because of punctuated equilibrium. Punctuated equilibrium (Eldredge and Gould 1972) helped us to look at the fossil record at a slightly different angle than we did previously, and with this new look, familiar patterns took on new meaning and novel phenomena became obvious possibilities. Of all the aspects of macroevolution, it was large-scale trends that benefited the most from punctuated equilibrium.

For decades before punctuated equilibrium, trends were not well understood. There were too many patterns to make sense of and there was no conceptual framework. Looking back, it seems like there was close to a one-to-one relationship between patterns and processes, for example “orthogenesis” the pattern and “orthoevolution” the theory, where every particular pattern was thought to be underpinned by a general process that produced it. The “law” of morphological evolution, for example Cope’s rule, Williston’s law (Gregory 1935), etc. are all of this time. G. G. Simpson (1944, 1953) succeeded in breaking this myopic view of trends by recognizing their statistical nature inherent in the bushy proliferation of species. His key insight was that patterns can vary over time and phylogeny without the need for a separate processes to explain all the variation. A single large-scale process could give rise to inconsistent smaller-scale patterns as different branches of a clade evolve independently and yet, still produce an overall trend.

Punctuated equilibrium further distilled Simpson’s statistical view of trends by identifying a single process that is flexible enough to produce most types of trends. Stanley’s (1975, 1979) idea of species selection is this type of process. It was first recognized because punctuated equilibrium suppresses any tendency for microevolution to underpin trends. The logic goes like this: if incessant microevolution did not in fact lead to sustained change within species (there is stasis instead), then there must be some more inclusive process that could shepherd populations of species and cause a trend. The shepherding process of species selection that Stanley identified maybe sufficient to produce many of the examples of trends we know from the fossil record because it can act in many directions and magnitudes. Species selection can act to produce trends when morphological stasis within species is common. It can move populations of species if there is no inherent bias in the phenotypic direction of speciation. It can even act when species evolve gradually and speciate directionally (McShea 2004, Simpson 2010b).

For Stanley, natural selection at the organismal level is more than a metaphor for species selection. For him the two levels of selection are isomorphic and species selection is as real a phenomenon as natural selection is. Selection among organisms and selection among species are essentially the same process, but they occur at hierarchically separate levels of organization. Populations of variable organisms, with the addition of natural selection, will show evolutionary change. And in just the same way, but with different players, populations of variable species with the addition of species selection results in macroevolutionary trends.

Not everyone has embraced species selection (e.g., Coyne and Orr 2004, Williams 1992). There are two hurdles to the acceptance of species selection, one philosophical and the other practical.

Many are philosophically opposed to the idea of species selection because of their belief that an understanding of all features of life and the physical world comes only by reducing everything to their lowest constituent parts. Although reductionism has been a powerful scientific tool, it is not the only one (Deutsch 2013). To me, this philosophical hurdle is easily passed without delving into philosophical argument, but rather can be shown to be false by using only paleobiological observations. Species selection has been empirically demonstrated in an increasing number of studies (e.g., Simpson and Harnik 2009; Goldberg et al. 2010; Mayrose et al. 2011; Powell and MacGregor 2011; Rabosky and McCune 2010; Simpson 2010, 2013). The irreducibility of speciation and extinction require us to reject a reductionist interpretation of these results.

The practical hurdle is more difficult, but it is also possible to pass. As scientists, we want to be certain that we can positively recognize species selection when it occurs and not be fooled some other phenomenon that can mimic it. A major step toward this goal came with the conceptual distinction between sorting and selection (Vrba and Gould 1986). Sorting’s utility came from its recognition of two sources of uncertainty; the uncertainty about the level at which selection acts in any given situation, and the uncertainty about the causes of selective patterns. At the time of Vrba and Gould’s paper, both uncertainties where rampant. Sorting provided a helpful way to think and discuss macroevolutionary patterns without devolving into an argument about the causes of those patterns before we had the tools to test those arguments. This was a big advantage at the time because few macroevolutionary patterns had been well documented. Sorting has traditionally lumped both sources of uncertainty together—any uncertainty about the level of selection or if differential diversification is caused or uncaused would classify a pattern as sorting.

My goal with this paper is to re-evaluate species sorting in light of current concepts, methods, and empirical results. I will argue that, we can be much more certain about the levels at which selection acts because of what we have learned about macroevolution over the last 30 years. For example, the demonstrable irreducibility of speciation and extinction shows that an emergent fitness view of levels of selection is correct. This means that any pattern of differential diversification is due to processes acting at the same level as diversification and not as a byproduct of selection at lower levels.

The concept of sorting also encompasses uncertainty in the type of causation of differential diversification, it recognizes differential diversification as an interesting pattern and does not distinguish between a caused selective pattern and a chance stochastic pattern. I think this source of uncertainty is conceptually simpler to understand if we do away with the concept of sorting and instead contrast the two macroevolutionary processes that underpin it: caused selection with non-caused drift. Drift in macroevolution, as it is within organismal populations, is an ephemeral and chance pattern of fitness that induces directional proliferation.

The gain by including species selection in our conceptual arsenal of macroevolution is at least as great as the ones that Simpson and punctuated equilibrium brought to trends. We can evaluate if species selection is real or not, as well as its relative frequency and importance in structuring macroevolutionary patterns, including patterns that I call non-trends, those patterns of major large-scale stability because of complex underlying dynamics where species selection and microevolution interact. One example of a non-trend is the stable frequency of coloniality and photosymbiosis in scleractinian corals (Simpson 2013). In this case, species selection and microevolutionary trait change act in concert to constrain the relative frequency of coloniality and photosymbiosis over 200 million years. Non-trends are phenomena that can only be recognized by studying species selection.

## Irreducible macroevolution

There are two known phenomena that are common in macroevolution but that are absent from microevolution—speciation and extinction. If speciation and extinction where absent, evolutionary patterns that take place over macroevolutionary time-scales would not be very different from the expectation of current neontological theory. In this counterfactual world, large-scale patterns would all be uniformitarian and result from the accumulation of the microevolutionary processes that we can observe directly today. But speciation and extinction actually happen. And as a consequence, the new dynamics characteristic of macroevolution become apparent. The macroevolutionary patterns we actually observe are not simply the accumulation of microevolutionary change because speciation and extinction introduce a threshold between micro- and macroevolution. This threshold arises because macroevolution occurs by changes within a *population of species*.

With every speciation event, a new independently evolving lineage is added. And with every extinction a species is removed. This may sound trivial, but these events are important because they have the ability to dominate the evolution within a population of species. Adding and subtracting species can change the distribution of traits in a population of species faster and in more ways than changes can be made with fixed species numbers.

## Speciation

Let’s look at speciation first. Speciation is the splitting of one or a set of interconnected populations into independently evolving populations. Speciation frees new species from demographic and genetic connections to other species, and allows the new species to follow their own path and evolve independently. This independence of mode among species directly affects all patterns of macroevolution that happen within whole populations of species. Independence makes it more difficult to explain large scale pattern as the cumulative effect of patterns happening within species.

Consider a hypothetical example, a single species of rabbit is evolving ever longer ears over time. After a million years, this rabbit species branches off a descendent species and both ancestor and descendent species coexist after the speciation event. The descendent, which is free to evolve independently from its ancestor, evolves shorter and shorter ears over time as its ancestor continues on its trajectory toward long ears length. The average ear length in the population of rabbit species will at first be a function only of the ancestral species and later, after speciation, the average ear length is a function of the two coexisting species (Fig. 1). In this example with the ear length of one species increasing and the other decreasing, the average does not change much after the speciation event as the net difference between both species increases as they diverge from each other in length.

**Figure 1:**
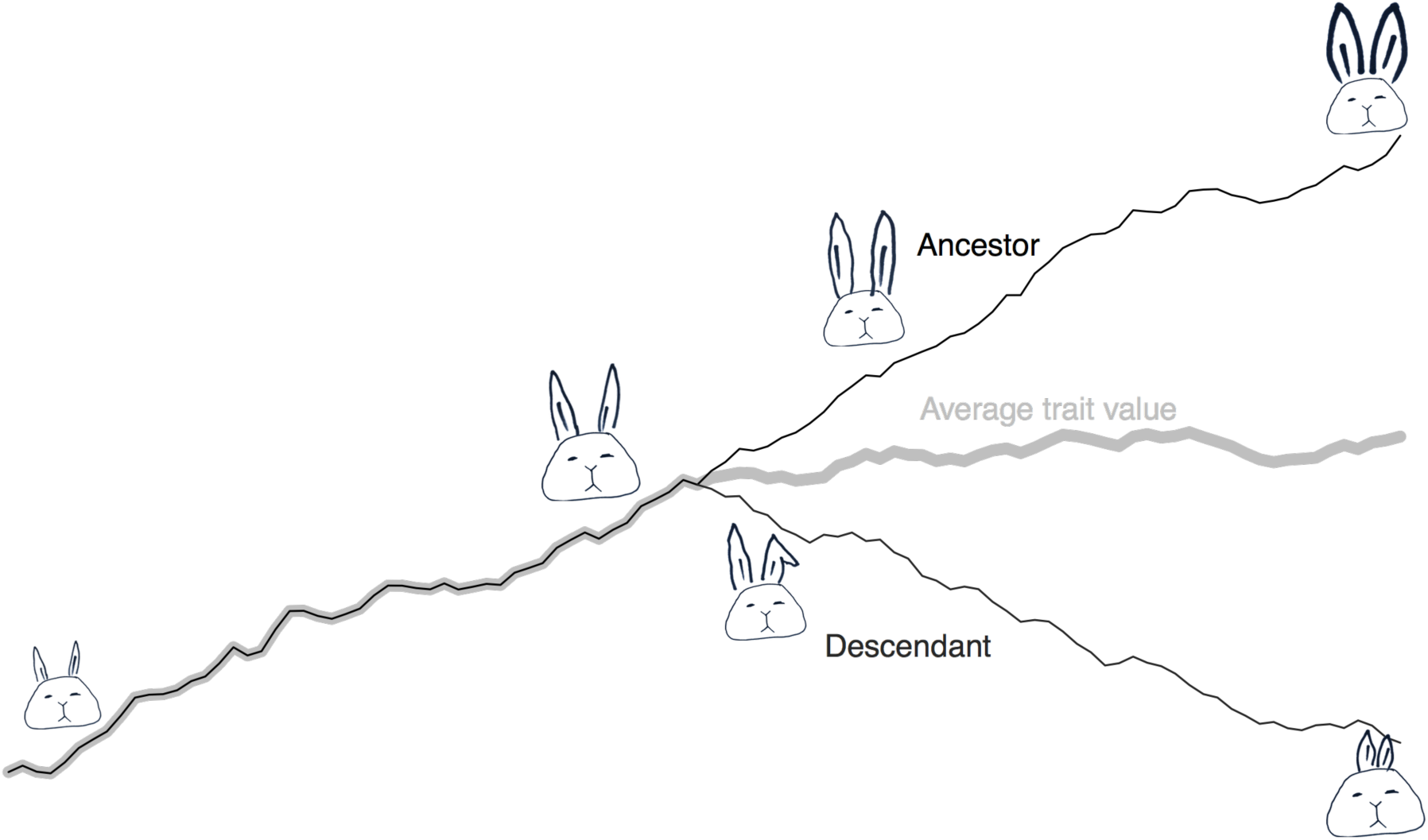
A cartoon of an anageneticly evolving rabbit species and its independently evolving daughter. The vertical axis shows rabbit ear length’ and time is shown on the horizontal axis and moves from left to right. The graph depicts the mean ear length of the population of species. When there is only one species, the mean ear length exactly tracks the single ancestral species until speciation. After speciation the mean incorporates the sizes of both independently evolving species. The more independent species there are in a population, the less the evolution of any one species contributes to changes in the overall mean among species. The lesson here is that for anagenesis to play a major part in a macroevolutionary trend, a significant proportion of the species within the population must show similar patterns of morphological change overtime— which is difficult when they are all evolving independently. This issue occurs in this simple twospecies example, but it gets ever more difficult to overcome as the species within a population become more numerous

Macroevolutionary trends are patterns that occur in populations of species, so any variation in evolutionary trajectory among species could easily cancel out the evolutionary change of others as the ancestor and descendent species of rabbits did to each other in the example. The more species are involved the stronger this averaging effect becomes. Only in a clade of one, is there is no averaging effect and only in this extremely rare case is the evolution of a species the same as the evolution of a clade.

## The causes of speciation rates

With each new species that evolves, not only is there the potential for the new species evolve independently from all the others. But there is also the potential for the rate of speciation to change with its introduction. Any variation in speciation rates within the species in a population has the potential to induce a trend.

Differential speciation (and extinction) constitute the raw material for species selection and sorting. Traits associated with species with relatively high speciation rates will increase in frequency over time. Simultaneously, traits associated with species that have relatively lower speciation rates will decrease in frequency. The pattern of covariation between rate variation and trait variation determines how the relative frequency of traits will change over time (Arnold and Fristrup 1982, Simpson 2010b, 2013, Simpson and Mller 2012), just as the covariance between fitness and trait variation among organisms determines how trait values will change by natural selection (Lande and Arnold 1983, Price 1970, Price 1972, Rice 2004).

Differential speciation can change the frequencies of any type of trait that it covaries with. It does not matter if the trait is emergent and inexpressible by single organisms, like geographic range size, or if the traits are attributes of individual organisms, like ear length. As long as there is a covariance between speciation rates and trait values, there will be a macroevolutionary change.

This ability for differential speciation to shepherd all types of traits, from organismal to emergent, has caused a lot of conceptual problems. Some researchers have restricted species selection only to the case where differential diversification directly covaries with emergent traits. Otherwise there is no way to tell if natural selection among organisms is the actual cause of the differential speciation. Vrba’s effect hypothesis (Vrba 1980, Vrba 1983) encapsulates this view, and states that any covariation between differential speciation and organismal traits must be caused by the effects of organismal-level natural selection percolating upward to the level of speciation. Restricting species selection to act only on emergent traits is thought by some to eliminate the possibility of confusing it with the effect hypothesis.

However there is a different school of thought that focuses on emergent fitness rather than emergent traits. This view recognizes a broader set of patterns as indicating species selection (Jablonski 2008). For this school, differential speciation is emergent fitness. For them it is important to note that the cause of differential speciation will always be at the species level. Traits at any level can covary with speciation rates and macroevolve, but organismal level traits do so by hitchhiking with a covarying intermediary trait.

Which of the two alternative conceptual frameworks, emergent traits or emergent fitness, is biologically most realistic? To test between these differences we need to know how differential speciation relates to differential organismal fitness. I see two mutually exclusive possibilities. First, differential speciation could be reducible to differential organismal fitness so that high speciation rates would be caused by high organismal fitness. Alternatively, organismal fitness and differential speciation could be independent features of life. In this case, the two things are hierarchically nested, within species, the constituent organisms have their fitnesses, while at the same time species have their speciation rates. The key to answering one way or the other is in identifying any link between the fitness of species (differential speciation rates) with the fitnesses of organisms within populations.

If there is a link between organismal fitness and speciation rates, then there will be a positive correlation between fitness and speciation rate. One way to identify such a correlation is to see if the temporal patterns of organismal fitness match the temporal patterns of speciation rates. If organismal fitness increases over time and these fitnesses are linked to speciation, then speciation rates should also increase over time.

It is not tractable to empirically measure organismal fitness over the lifetime of a species. Instead, we must turn to theoretical expectations on the temporal pattern of fitness (e.g., Orr 2009). There has been long argument within the microevolutionary community about how fitness changes over time. Wright, claimed that average fitness always increases in proportion to the slope of the fitness landscape. As populations climb the hills of the fitness landscape, they follow the steepest slope. And even as the landscape changes over time, the population will always take the steepest path upslope, and therefore always increases over time. Fisher (Fisher 1930) argued that there is more to the change in fitness than the slope of the fitness landscape. He argues that the rate of fitness increase is also proportional to the additive genetic variance, a pattern known as Fisher’s fundamental theorem. This means that traits with more variation will evolve faster than those with less variation. This structure of the genetic variance combines with the slope of the fitness landscape to determine the rate of fitness increase. In Fisher’s case variances are always zero or positive, so it converges with Wright’s view that the first order long-term expectation of fitness change is that it will increase over time, although how quickly the population climbs is modulated by both the variance in Fisher’s view and modulated by the change in the fitness landscape in Wright’s view.

Since Fisher’s time we have developed more theory detailing the importance of the specific structure of genetic variation. Certain genetic interactions like epistasis and linkage disequilibrium, as well as population interactions like frequency dependence, can lead to a temporary decline in fitness. Even if these declines in fitness occur in specific genotypes within the species’ population, the mean fitness of all members of a species will likely increase as genotypes replace each other successionally over time. So the long-term trajectory of organismal fitness should follow Fisher’s and Wright’s expectations.

If speciation rate is linked to organismal fitness by upward causation (as expected given the effect hypothesis) it too should increase over the lifetime of the species. The easiest way to empirically measure this temporal pattern in speciation rate would be to tally the frequencies of speciation events over a species’ lifetime. Although this has rarely been been done, two recent studies have looked into the relative timing of speciation in the fossil record (Liow and Ergon 2015) and with molecular phylogenetics (Hagen et al. 2015). With the few examples of species-level phylogenies including explicit ancestors available in the fossil record Liow and Ergon (2015) fit models of age-dependent speciation (including the frequency of speciation given a temporal duration) and identify several different patterns of age-dependence. In some groups biased toward the early part of the ancestor’s lifetime, and others they find that long-lived species have higher speciation rates. Using molecular phylogenies of extant species, Hagen et al. (2015) found that the frequency of speciation is ubiquitously biased toward the beginning of an ancestor’s lifetime—a result counter to the expectation of a link between organismal fitness and speciation rates.

Here are a few additional examples that are useful in identifying how the pattern of speciation over the lifetime of ancestral species can vary. One example, planktonic macroporiferate foraminiferans (Aze et al. 2011) show a uniform patterning of speciation along the lifetime of ancestors (Fig. 2)—speciation is as likely to occur early in an ancestor’s lifetime as it is toward the end (similar to the pattern Liow and Ergon 2015 found). The phylogeny links ancestor-descendant pairs of morphospecies together and records the timing of first and last appearance of each morphospecies in the geological record. Here, I measured when, relative to the ancestor’s lifetime, speciation occurred. The timing of speciation in the cheilostome bryozoan *Metrarabdotos* (Cheetham 1986, Cheetham et al. 2007) provides a second example. In *Metrorabdotos* the timing of speciation is clustered in the first half of species lifetimes (Fig. 2). This result is different than the old-age dependent speciation Liow and Ergon find for *Metrarabdotos*, because they identify a tendency for longer-lived species to have more descendants independent of when those descendant species were budded off. Lastly, speciation in the trilobite genus Phacops (Eldredge 1971, 1972) is clustered early in the lifetime of ancestors (Fig. 2).

**Figure 2:**
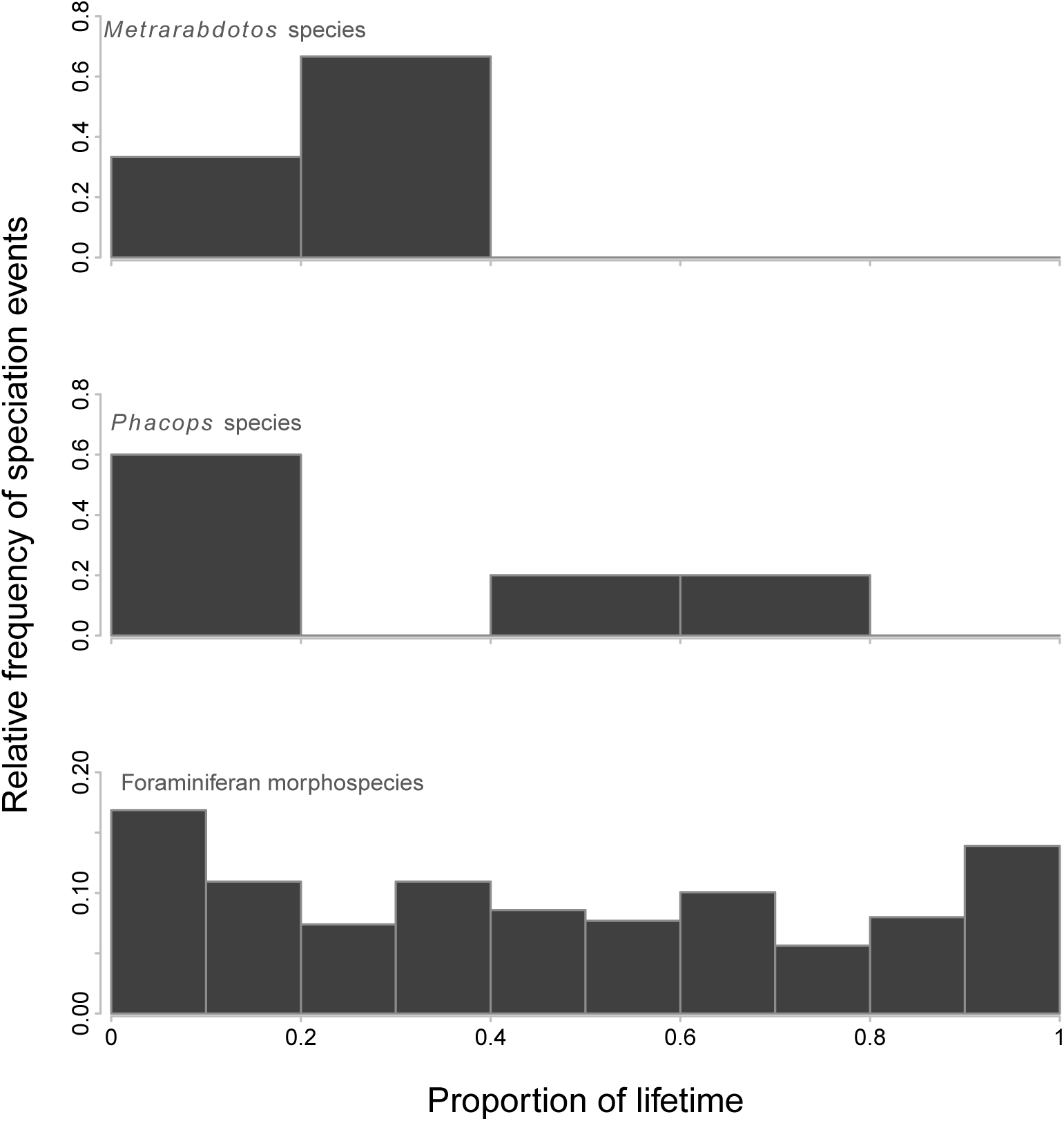
A tabulation of the relative frequency of speciation events during the relative age (lifetime) of ancestors. The three groups of fossils shown here have are based on phylogenies with explicit ancestor-descendent relationships defined. The top panel shows the relative frequency of the timing of speciation events within the cheilostome bryozoan *Metrarabdotos* (Cheetham et al. 2007). The middle panel shows the relative frequency of the timing of speciation in the trilobite *Phacops* (Eldredge 1972). The bottom panel shows the relative frequency of the timing of speciation for species of macroperforate planktonic foraminifera (Aze et al. 2011).

These three examples, as well as the patterns seen in Liow and Ergon (2015) and Hagen et al (2015) reject the possibility that speciation is reducible to organismal fitness. Like cell-division and sexual reproduction, organismal fitness and speciation are independent processes that occur together in a hierarchically nested arrangement of organisms within species. The two levels of fitness may interact, but they are independent, and caused independently.

## The causes of extinction

Compared to the irreducibility of speciation, extinction seems, at first glance, to be easily reducible. After all, a species of must be extinct if all its constituent organisms are dead. Nevertheless, this line of thought leads to the false conclusion that species go extinct due to the same causes from which organisms die. In the end our conclusions about the reducibility of extinction should focus on distinguishing its causes compared to those that cause organismal fitness.

The causal link between death and extinction is tenuous. Organisms die from the numerous things that can kill them; it could be predation, disease, starvation, accident, the weather, fights, or old age that does the deed for any particular organism. The cause of death will always vary from organism to organism within a population. The net effect of all this death is that the population may shrink if the birth rate is not sufficiently high to replace the numbers that are killed. Population decline maybe a common cause of the extinction, but organisms never die from population decline, and population decline is never caused by organismal death alone because birth rate plays a major role in population growth. It’s the relative balance between birth, death, and lifespan of organisms that determines population growth rate and consequently extinction.

Moreover, extinction can be guaranteed for a species prior to the collapse of its populations, as is the case when populations are below a minimum viable population size. In this situation extinction is caused by the population crossing a threshold of viability. Organisms within the population do not, however, die from being a part of an inviable population. Instead, they succumb for any number of more mundane reasons. The Pinta Island tortoises went extinction for a very different reason than the heart failure that killed Lonesome George.

There can be a positive correlation between organismal fitness and population growth rates. This correlation, termed the Allee effect, is particularly important in frequency dependent and limited populations. The existence of this effect is likely an example of downward causation, where the fitness of organisms are influence by their population context.

Two paleobiological patterns suggest other causes of extinction that don’t percolate up from organismal fitness. The first, termed Van Valen’s law (or the law of constant extinction) is a characteristic pattern of extinction that emerges among genera or families within a higher taxon (Raup 1975, Van Valen 1973). Van Valen identified a distinctive log-linear pattern in the age-frequency distributions of the majority of fossil higher taxa. The log-linearity in this distribution means that, within a higher taxon, genera (or families) of all ages share the same probability of extinction. A long lifetime does nothing to buffer genera or families from extinction. Van Valen was quite surprised with this result because natural selection among organisms is expected to increase their fitness (and consequently their adaptedness) over time as their populations evolve. If true, Van Valen’s law shows that extinction is decoupled from the patterns of organismal fitness. Van Valen proposed the Red Queen’s Hypothesis to give a possible mechanism for this decoupling by invoking the importance of competitive interactions among species, genera, or families within an adaptive zone. The biotic environment is so shifty and complex that even ever increasing fitness within a species does not necessarily increase higher fitness relative to other competing species.

The second paleobiological pattern is the symmetric rise and fall of species and genera (Foote 2007, Foote et al. 2007, Jernvall and Fortelius 2004, Liow et al. 2010, Liow and Stenseth 2007). On average, the occupancy or range size of species rises from a low initial value to a peak halfway through the lifetime and subsequently falls until extinction occurs. As with the age-independence of extinction, the rise-and-fall pattern cannot be explained by changes in organismal fitness because those are expected to increase over the lifetime of a species. The cause of the rise and fall pattern is not currently known. From interval to interval there is a correlation between the second order changes in occupancy and a change in environments (Foote 2014). But I think that this environmental cause is unlikely generalize to the first-order rise-and-fall pattern that plays out over lifetimes, because any time the environment changes in one direction, species of young and old ages are both affected. When species are lined up from birth to death, rather than from the time in the past they occur, the effect of any specific event of environmental change that is experience by young and old alike will be smeared across the lifetime and on average cancels out. As with the Red Queen, biotic interactions among species may be critical to understand the rise-and-fall pattern (Simpson 2010a).

As we have seen, extinction is a clearly emergent phenomenon. The features it has are not the same as the features of traits that are ambiguously emergent. Often in ambiguous cases, variability plays a part in the irreducibility of a phenomenon consisting of traits that are reducible. For example, body size is an organismal trait, yet because it is relatively invariant within a species, it can also behave similar to an emergent trait—the species has a characteristic body size that can be selective (Van Valen 1975). Extinction is emergent for the opposite reason and in spite of the variable causes of death of its constituents.

Differential extinction can change the frequencies of traits over time by culling species away. It can also act in conjunction with differential speciation through selective net diversification. Most of the patterns of selectivity that speciation can produce occurs inversely with extinction. Unlike speciation, extinction’s power to create trends is immediate because the loss of species immediately culls the trait distribution.

## The many levels of selection

The philosophical issue that prevents the easy recognition of species selection hinges on a suspicion that species selection is reducible to natural selection. Any confusion about the level at which selection acts in any given scenario would get in the way of a useful theory.

The concept of species sorting incorporates this uncertainty about the level of selection by being agnostic about the level that differential speciation and extinction rates are caused. It recognizes and highlights the pattern of differential diversification without concern about the level of selection that causes the pattern. But the fact that speciation and extinction are emergent removes this uncertainty about the level at which selection acts. Any pattern of differential speciation, extinction, and diversification are themselves emergent because these differentials are created by emergent patterns of speciation and extinction and not caused by organismal fitness. Speciation and extinction are the macroevolutionary analogues of birth and death in organismal fitness, and as such constitute an emergent species-level fitness. Organismal and species level fitnesses occur together in much the same way that zooid and colony level fitnesses both occur in colonial organisms (Simpson 2011, 2012). Fitness represents a demographic process at each level that it occurs. As a consequence, any trait that covaries with diversification rates has the potential to change by species selection.

This certainty in the level at which species selection acts—at the species level due to emergent and differential diversification—helps us make the next step and solve the practical problem, which is how to recognize species selection when it occurs.

## Selection and drift—the caused and the uncaused

The concept of species sorting incorporates another source of uncertainty about differential diversification—uncertainty about how the rate differentials are caused. There are two ways to be uncertain about these causes. First, are the traits associated with the rate differential acting to directly cause the rates or are they a part of a chain of causation that links the traits we measure to diversification, essentially hitchhiking with some unmeasured intermediaries? And second, are the differential rate patterns caused at all or are they temporary random associations and therefore uncaused. Hitchhiking is likely to be extremely common in nature. Most species are likely to have a number of emergent traits and a significant number of organismal traits that covary with each other. Selection in a multivariate regime can be quite complex with characters evolving due to their interactions with each other and with fitness. Even if a trait is evolving only by hitchhiking through a covariation with the trait that is the target of selection, it can effect the outcome of the selective process. The resulting evolutionary change will deviate from the trajectory that would have occurred if the target of selection was able to vary independently with the hitchhiking trait. In other words, the structure of covariation between selection and many evolving traits is important in determining the response to selection (Lande and Arnold 1983, Rice 2004). Chance associations between traits and fitness are also common in micro-evolution and known as drift. Drift mimics natural selection and it can easily lead to evolutionary changes in small populations, where common stochastic differences in fitness between organisms can take on a large effect. Drift is the same at the species level. With the emergent fitness approach, selection, whether direct or via hitchhiking, and drift are the only two processes encompassed by species sorting. Selection and drift are real evolutionary processes but sorting is a conceptualization of our scientific knowledge. As such, sorting can be a useful tool to keep us asking the right questions and organizing our uncertainty. But only selection and drift are evolutionary processes. If we recognize both processes of selection and drift, then the concept of sorting becomes a redundant and imprecise conception of actual macro-evolutionary processes.

## Conclusions

We know that life has hierarchical structure that has a physical manifestation and also an expression in processes like fitness. The physical manifestation is simply cells within bodies, bodies within populations, populations within species, and species within populations of species. The expression of hierarchical structure in terms of fitness occurs because the parts and wholes contained in that list can be added by birth or speciation, subtracted by death or extinction, or expand in size by the births and death of constituent parts. The physical manifestations are what does the evolving where as the demographic changes expressed by fitness are involved in causing that evolution.

To understand how to distinguish between selection and drift as well as understand the causal structure that associates traits and fitness, we need to have a precise way to calculate patterns of selection and microevolution. I’ll provide a framework to do this in a companion paper (Simpson, Species selection on evolving characters). I think that this framework will help us to understand how all the processes that we know about interact with each other. With that knowledge, we can then focus in real-world situations and apply this macroevolutionary framework to understand what’s going on. My hope is that this framework will give us a way to use mismatches between empirical observations and theoretical predictions as well as the other surprising results to enrich our knowledge of macroevolution. For too long, the field has used empirical observations as evidence for or against two fixed and opposing conceptual positions. With this framework I hope that empirical observations will be used to extend past these two conceptual positions develop and refine our understanding of macroevolution.

## Acknowledgments

Thanks to Doug Erwin, Dan McShea, Chris Haufe, Sarah Tweedt, Rachel Warnock, and Jenna Rolle for discussion and comments. This paper was born from listening to Warren Allmon’s questions to my talk during the Florida NAPC in 2014. I realized that certain parts of macroevolutionary theory have developed quickly over the years but that many of those ideas and arguments have not received enough attention in print. It’s time to get them out of the shadows so we can see if they are worth developing further.

